# Direct estimation of mutations in great apes reveals significant recent human slowdown in the yearly mutation rate

**DOI:** 10.1101/287821

**Authors:** Søren Besenbacher, Christina Hvilsom, Tomas Marques-Bonet, Thomas Mailund, Mikkel Heide Schierup

## Abstract

The human mutation rate per generation estimated from trio sequencing has revealed an almost linear relationship with the age of the father and the age of the mother. The yearly trio-based mutation rate estimate of ~0.43×10^−9^ is markedly lower than prior indirect estimates of ~1×10^−9^ per year from phylogenetic comparisons of the great apes. This suggests either a slowdown over the past 10 million years or an inaccurate interpretation of the fossil record. Here we use sequencing of chimpanzee, gorilla and orangutan trios and find that each species has higher estimated mutation rates per year by factors of 1.67+/− 0.22, 1.54+/− 0.2 and 1.84+/− 0.19, respectively. These estimates suggest a very recent and appreciable slowdown in human mutation rate, and, if extrapolated over the great apes phylogeny, yields divergence estimates much more in line with the fossil record and the biogeography.

## Introduction

The current human mutation rate has been extensively studied through sequencing of thousands of trios (*1–3*). The consensus is that the rate increases almost linearly with the age of the parents with a higher rate of mutation from the father (2.51 per year) than from the mother (0.78 per year), yielding a mutation rate of 0.43×10^−9^ per base pair per year (*2*). The mutation rate is important for calibration of the molecular clock for dating evolutionary events in human ancestry, such as human population divergences, human-Neanderthal divergence and the split time with ancestors of the other great apes such as chimpanzees, gorillas and orangutans (*4–6*). Extrapolating the present yearly human rate estimated from trios predicts average genomic split times with chimpanzees of more than 15 million years and orangutans at 35 million years and these estimates are difficult to reconcile with the fossil record (*5, 7–10*). It is possible that the mutation rate could have decreased over time in the lineage leading to humans. However, the great apes phylogeny almost supports a molecular clock with the chimpanzee, gorilla, and orangutan branches only being 2-3%, 6-7% and 11% longer than the human branch (*7*). Thus, a decrease in human mutation rate per year would therefore have to be either very recent, be of small magnitude, or there might have been independent decreases in the different great apes lineages, perhaps due to a general longer generation time in all lineages (*7, 11, 12*). One way to distinguish among these alternatives is to determine the present mutation rate in other great apes from trios. The first such study using a pedigree with six chimpanzee trios (*13*) reported a very similar per generation rate as in humans and a more male biased mutation contribution. However, the trios used had young parents suggesting a higher yearly rate than in humans. A recent study using deeper sequencing suggests a higher rate than in humans by about 30%(*14*).

## Results

Here we call mutations in extended trios for chimpanzees, gorillas and orangutans sequenced to high coverage and combine our results with a reanalysis of the raw data from Venn et al (*13*). We called *de novo* mutations using stringent quality threshold and used a probabilistic model to estimate the fraction of the genome where we would call *de novo* mutations—if they were there (*15*). The mutation rate is the fraction of these two quantities and the quality threshold was determined as the minimum quality value where increasing the quality does not change the estimated rate (Q60, see supplementary Figure S1). The number of mutations called for each of the three species, together with our reanalysis of the previous published sequencing data set from six chimpanzee trios of an extended pedigree (*13*) is shown in Figure 1. The estimated rates take the number of callable sites into account (presented in Supplemental Table S2) and is shown for each trio in Figure 1. The number of mutations that could be assigned to parental origin using read-backed phasing are also shown. The distributions of mutations along the genome in the trios show a slight clustering as reported from humans (supplementary Figure S2) and there are no significant differences in their composition (supplementary Figure S3) among species. Furthermore, the estimated rates are very similar for the repetitive and non-repetitive parts of the reference genomes (supplementary Figure S4), and the number of transmitted mutations to grandchild is not significantly different from 50% (supplementary Figure S5). The estimated mutation rate correlates with paternal age across species (Figure 2a).

**Figure 1.**
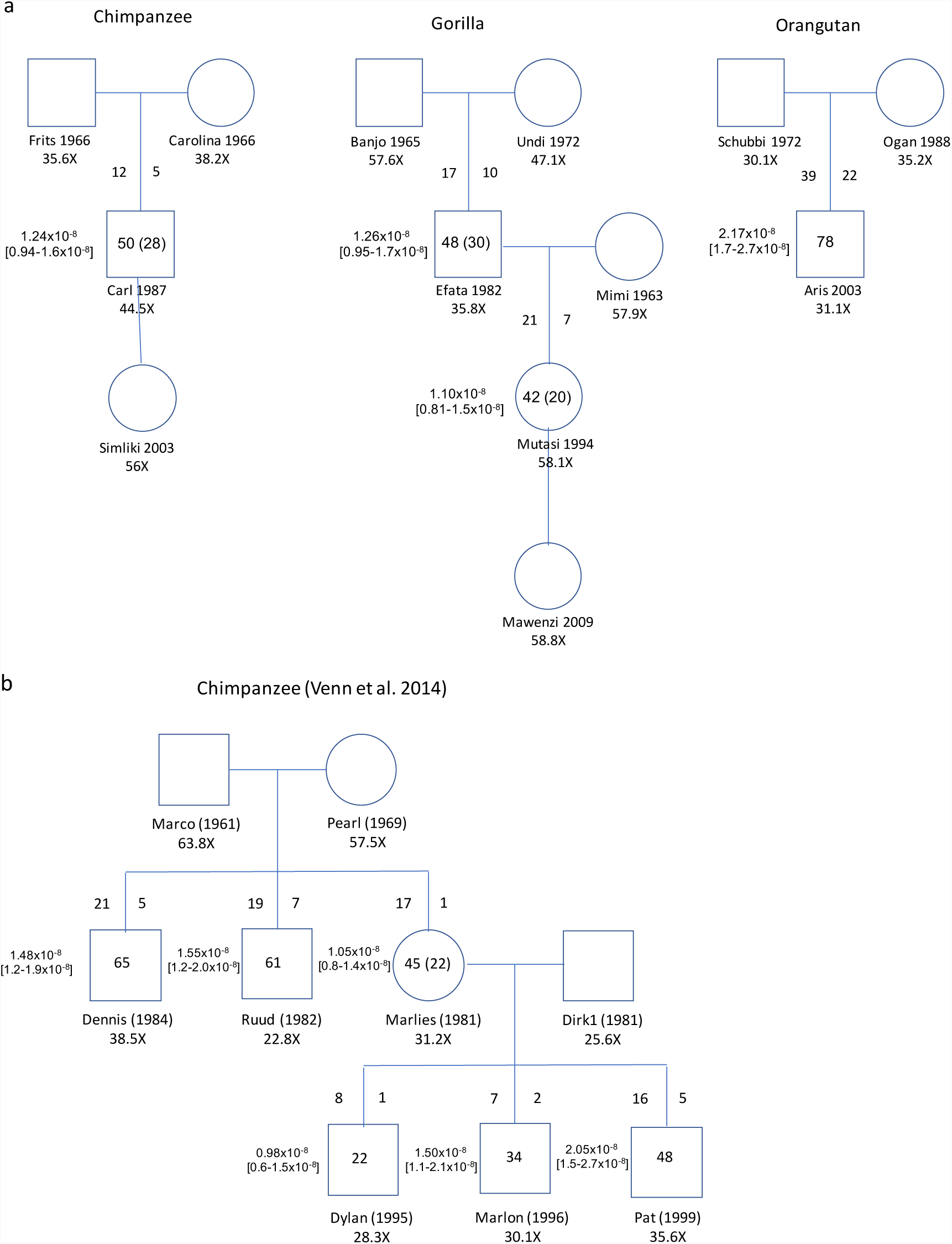
Numbers, rates and transmission of *de novo* mutations. a) The sequenced pedigrees of chimpanzee, gorilla and orangutan with names, birth years, and sequence coverage. The number of mutations observed within individuals and the number of transmitted mutations in parenthesis. On the left to the transmission line is the number of mutations inferred to have arisen in the father, on the right the number of mutations inferred to have arisen in the mother. Next to the individuals is the inferred rate [95% CI]. b) Results from the reanalysis of the chimpanzee pedigree sequenced by Venn et al.

**Figure 2.**
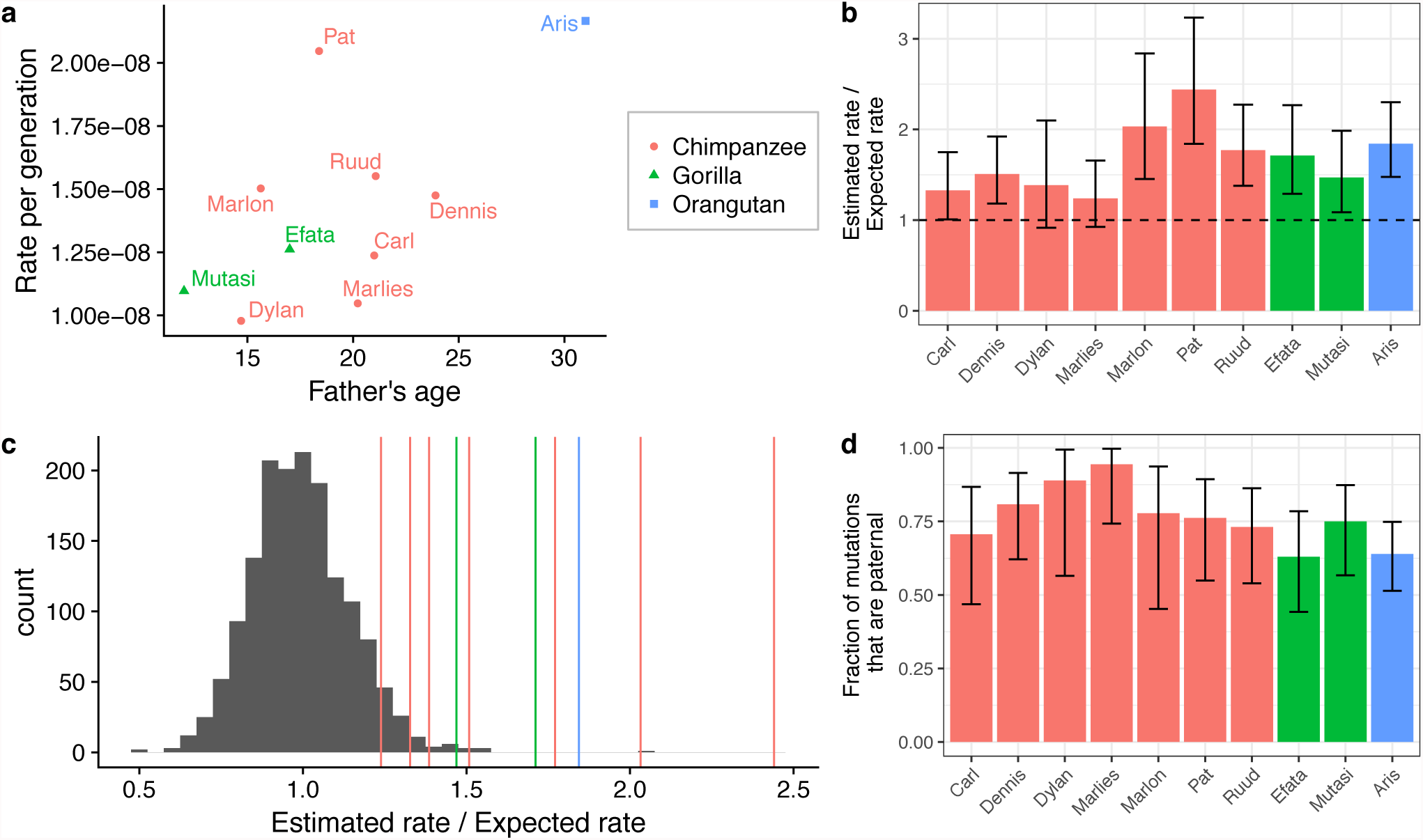
Properties of *de novo* mutations. a) The estimated mutation rate as a function of paternal age. b) The estimated rates compared to the expected rates (95% CI). c) The histogram shows the distribution of Estimated / Expected rates for the human samples from (*2*), and the vertical lines show the Estimated / Expected rates for the analysed great ape trios. d) The fraction of the mutations for which parent-of-origin could be assigned that came from the father (95% CI).

In order to turn the observed mutation rates in single generation pedigrees into a yearly mutation rate estimate for the different great apes species, we extrapolated the relationship with maternal and paternal age observed in humans (*2*) to the other great ape species. This relationship is: Mutation rate = 1.77×10^−9^ + 7.26×10^−11^ * maternal age (years) + 2.87×10^−10^ * paternal age (years). This provides an estimate of the mutation rate expected for each offspring in the trios from the age of its parents at the time of birth. This expected rate is then compared to the observed rate in order to test for a change in yearly rates among the different great apes species (see Table 1). Figure 2b and Table 1 shows that under this strong assumption, the estimated yearly rates for chimpanzee, gorilla and orangutan are each significantly higher than in humans, but not significantly different from each other. Figure 2c shows these ratios compared to the range observed in 1548 human trios (*2*), this is shown as a function of paternal age in Supplemental Figure S6.

**Table 1.** Basic statistics for *de novo* mutation calling

| Child | Species | Father age | Mother age | Callable sites | Total mut | CpG | Non-CpG strong | Weak | Rate observed | Rate expected | Relative rate |
| --- | --- | --- | --- | --- | --- | --- | --- | --- | --- | --- | --- |
| Carl | Chimp | 21 | 21 | 4041209937 | 50 | 9 | 24 | 17 | 1.24E-08 | 9.32E-09 | 1.33 |
| Pat | Chimp | 18.39 | 18.48 | 2345308475 | 48 | 3 | 25 | 20 | 2.05E-08 | 8.39E-09 | 2.44 |
| Dennis | Chimp | 23.9 | 15.89 | 4406543730 | 65 | 12 | 30 | 23 | 1.48E-08 | 9.78E-09 | 1.51 |
| Ruud | Chimp | 21.07 | 13.06 | 3931520576 | 61 | 20 | 22 | 19 | 1.55E-08 | 8.76E-09 | 1.77 |
| Marlies | Chimp | 20.22 | 12.22 | 4295124822 | 45 | 9 | 22 | 14 | 1.05E-08 | 8.46E-09 | 1.24 |
| Dylan | Chimp | 14.7 | 14.79 | 2248382773 | 22 | 6 | 12 | 4 | 9.78E-09 | 7.06E-09 | 1.39 |
| Marlon | Chimp | 15.63 | 15.72 | 2262663575 | 34 | 8 | 11 | 15 | 1.50E-08 | 7.39E-09 | 2.03 |
| Efata | Gorilla | 17 | 10 | 3804369957 | 48 | 7 | 20 | 21 | 1.26E-08 | 7.37E-09 | 1.71 |
| Mutasi | Gorilla | 12 | 31 | 3830238966 | 42 | 8 | 20 | 14 | 1.10E-08 | 7.46E-09 | 1.47 |
| Aris | Orang. | 31 | 15 | 3600614878 | 78 | 19 | 25 | 34 | 2.17E-08 | 1.18E-08 | 1.84 |

Due to a higher number of cell divisions in the male germ line than in the female germ line, the mutation rate in humans is male biased. In all three great ape species we also find the majority of mutations passed on from the father, with the male bias highest in chimpanzee as reported previously (*13*), but the differences among the species are not statistically significant (Figure 2d, Table 1). Among the mutations that can be assigned to parental origin (50-80% for the different trios), the number of mutations passed on increase significantly with paternal age but not with maternal age when species are combined (Supplemental Figure S7).

With these new mutation rate estimates we can revisit dating of the genomic divergence of the great apes. The great apes phylogeny is very close to adhering to a molecular clock with the chimpanzee branch slightly longer than the human branch (by 2%) and the gorilla branch longer by 6% (*7*). For this to be consistent with our estimated rates, a slowdown in mutation rate in the human lineage must be very recent. It is therefore possible that even the dating of the human Neanderthal divergence is affected. This is in accordance with the higher rates by 20% observed when using the recombination clock (*6*) and also the branch shortening estimates from using the 45,000 year old human fossil (*16*).

Figure 3 shows the timing estimates of great apes divergence based on extrapolation of the human rate estimates (green), and based on extrapolating the estimates we have obtained on the respective branches consistent with reported deviations from the molecular clock (Figure 3 (blue), see also Methods for details). To turn these divergence time estimates into species separation time estimates we used previous estimates of ancestral effective population sizes(*10*), calibrated to the new rates (red numbers in Figure 3).

**Figure 3.**
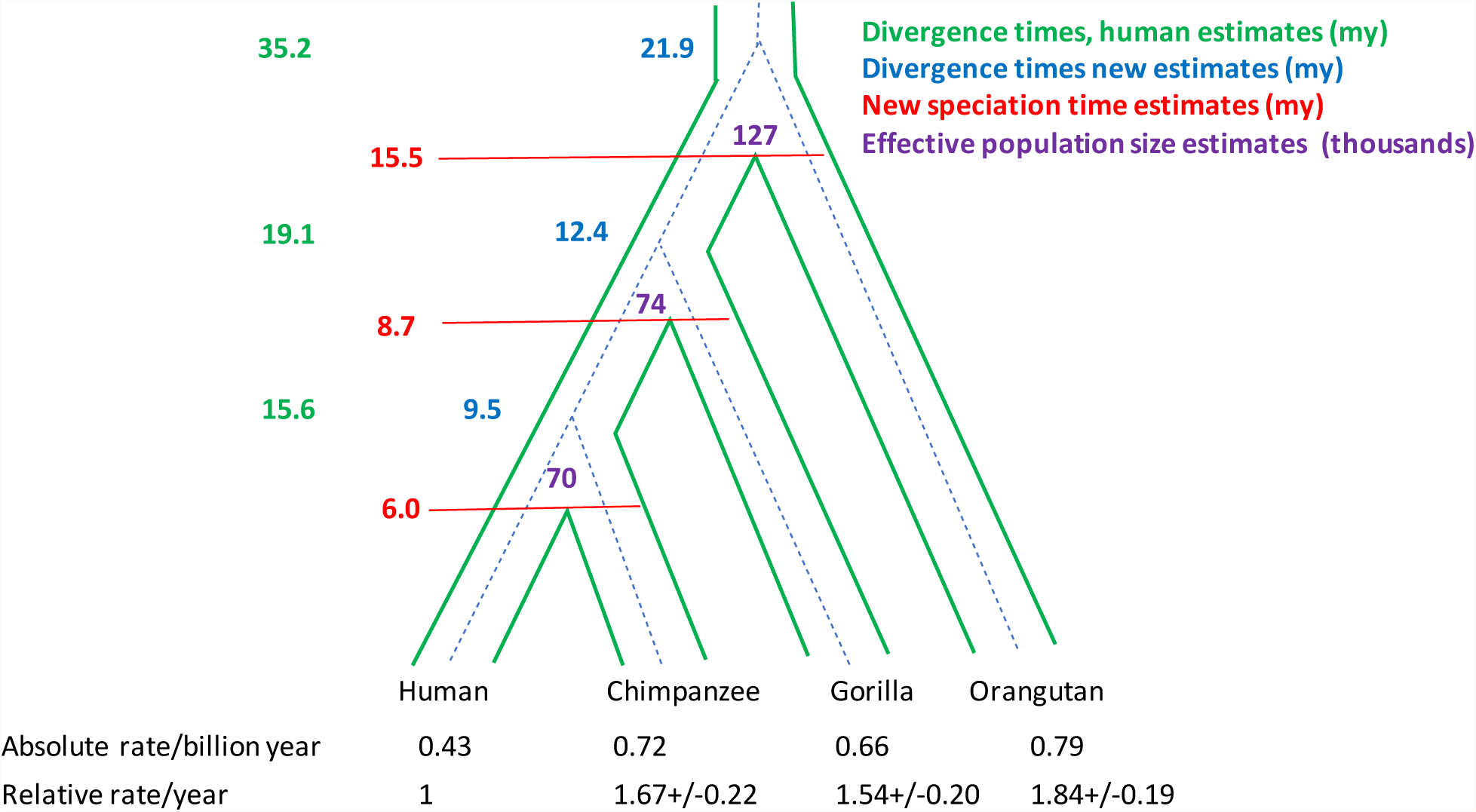
Estimates of great apes genomic divergence time and species separation times. The estimated mutation rates per year for each species is extrapolated over the phylogeny to yield divergence time estimates (blue numbers) which are more recent than estimates based solely on the current estimate of the human mutation rate (green numbers). Estimated ancestral effective population sizes (pink numbers, in thousands) are used to estimate speciation separation time (speciation time, red numbers). Absolute and relative rates per year (+/− SD) calculated from human rate extrapolation, combined over individuals for each species. See supplement for details

The apparent slowdown in mutation rate in the human lineage may be partly explained by humans having later onset of puberty and a longer generation time (human estimate 29 years, chimpanzee 24 years, gorilla 19 years, orangutan 25 years) (*2, 11*). Later onset of puberty implies more years with very few mutations accumulating in males. However, while these life history characteristics can explain the differences in branch lengths in the great apes phylogeny (*11*), they do not explain a reduction in mutation rate by about 40% in human trios compared to great apes trios. Most trios investigated have young parents (average 16.7 years females, 19.5 years males). This could explain a higher rate if the rate grows sub-linearly with parental age which, however, is not supported from human data (*2*). Estimates of human mutation rate further back in time using the recombination clock have been reported as 0.55+/−0.05 per bp per billion years (*6*). This rate falls almost exactly in between the trio-based rate in humans today and our estimated rates. It is possible that the large part of the slowdown in humans is very recent and perhaps caused by life style or environmental differences in the predominantly Caucasian populations where trios have been investigated for *de novo* mutations.

The speciation time estimates of great apes based on the new mutation rate estimates assuming a human slowdown pushes all speciation events closer to present time. A human chimpanzee separation time of 6.0 million years as we estimate is consistent with *Ardipithecus* (*17*) being on the human line, *Orrorin* (5-6 my old, (*18*)) being very early on the human lineage and *Sahelanthropus* (6-7 my old) being right at the split between human and chimpanzee. A species separation time of 15.5 mya with orangutan is well within the time interval between the age of *Sivapithecus* (12.2 mya, (*19*)) and *Proconsul* (23 mya), generally assumed to be lower and upper bounds for human orangutan divergence.

## Materials and Methods

### Samples

All blood samples were taken during routine health checks, and Convention on International Trade in Endangered Species of Wild Fauna and Flora (CITES) permits were obtained from countries outside the EU.

### Sequencing

Genomic DNA was extracted directly from EDTA whole blood samples using a DNeasy Blood and Tissue Kit (Qiagen, Valencia, CA, USA), following the manufacturer’s instructions. 2 ug DNA was used for construction of PCR free libraries with an average insert size of 250 bp. The libraries were sequenced on Illumina HiSeq X instruments by Novogene using standard chemistry for paired end sequencing of 2×150 bp to coverage between 30X and 56X (See Figure 1).

### Reanalysis of chimpanzee data of Venn et al

We downloaded the fastQ files from ftp://ftp.well.ox.ac.uk/panPed and analysed alongside our own sequencing data using the same pipelines.

### Mapping

Reads were aligned to the following great apes reference genomes, respectively;

Chimpanzee Pan_tro 3.0 (UCSC: panTro5)

Gorilla GorGor4.1 (UCSC: gorGor4)

Orangutan WUGSC2.0.2 (UCSC: ponAbe2)

Using BWA version 0.7.15. The average mapped coverage of the new data is shown in Figure 1 and Supplemental Table S1, and 90.8% of the aligned genome covered by at least 15 reads

### Variant calling

The realigned and Base Quality Score Recalibrated BAM files were used as input for multi-sample genotyping for each species separately using the HaplotypeCaller of the Genome Analysis Toolkit version 3.8.

### Identification of *de novo* single nucleotide variation

To limit the number of false positives we only consider a Mendelian violation as a possible *de novo* mutation if both parents in the family in question are homozygotes for the reference allele and if the variant is not called in any of the other families.

We apply these filters when we look for *de novo* mutations:

1. *A site filter* that look at the reads from all individuals to filter away bad sites that are not true variants. The site filter uses the following parameters:

a. FS: Fisher’s exact test on strand bias.
b. ReadPosRankSum: Rank sum test on position of the alternative allele in the reads.
c. BaseQualityRankSum
d. MappingQualityRankSum
2. *Individual filters* that look at the reads and genotype call of a single individual to discard a possible *de novo* call if we are not sure that all of the individuals in the family in question are called correctly. We use two different kinds of *Individual filters*:

a. *Homozygote reference filter*. This filter is applied to the parents to check that we have confidence that they really are homozygous for the reference allele. The filter uses the following parameters:

i. GQ: Genotype quality of the individual.
ii. DP: number of reads for this individuals at this site.
iii. AD2: number of times the alternative allele is seen in this individual.
iv. lowQ_AD2: the number of low quality reads (not used in the calling) that contained the alternative allele.
v. RC: the number of reads in this individual relative to the average coverage of this individual
b. *Heterozygote filter*: This filter is applied to the child to ensure that the child really is heterozygous at this site. The filter uses the following parameters:

i. GQ: Genotype quality of the individual.
ii. DP: number of reads for this individuals at this site.
iii. AlleleBalance: The fraction of the reads in the individual that contains the alternative allele.
iv. minStrandCount: The minimum number of counts of the alternative variant on each strand
v. RC: the number of reads in this individual relative to the average coverage of this individual

### Estimating callable sites for *de novo* mutations

We calculate the denominator of the rate estimate following the same strategy as also previously described in Besenbacher et al.(*15*)

To achieve a better estimate of the rate of *de novo* mutations in a trio, we base the denominator of the rate estimate on the probability at each site that we can call de novo mutation rather than simply counting a site as either callable or non-callable. The probability of calling site *x* as a *de novo* mutation given that it is a true *de novo* mutation in the family *f* we name the callability and denote it by 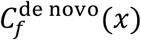. The callability can be estimated independently for each family based on the depth of the family members at the site, and the expected number of callable sites in a given family is then the sum of the callability of all sites in that family.

Since the site filter is based on statistical tests that follow a known distribution we can estimate how many good sites we expect to be filtered away by this filter by looking at the null distribution of the tests and assuming that the two tests are independent. We denote by *α*_*site*_ the fraction of good sites that we expect to be filtered away.

The mutation rate of a family *f* can then be estimated as:

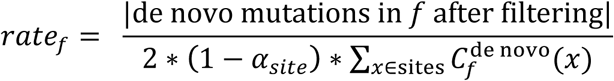

Now let *Z* be a genotype (Hetero, HomRef or HomAlt) and consider for an individual *i* the probability of calling it as *Z* at position *x* (and not filtering it away) given that the individual truly is *Z* at *x*. We denote this conditional probability by 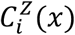 and it signifies the ability to give a true call of *Z* at *x*. Clearly this will be a function of sequencing quality at *x* (not least the depth). If we assume that the ability to truly call each member of a family is independent, then the callability of a site in a given family can be calculated as the probability of calling each individual correctly after filtering:

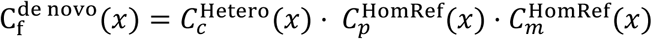

Where *c*, *p* and *m* indicate the child, father and mother of the family *f*.

Assuming that 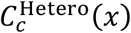 is independent of the parental genotypes as long as they are conducive to a heterozygous offspring, we can estimate it by considering only variants where one parent is homozygous reference with high confidence and the other parent is homozygous for the alternative allele. At such sites the child should always be a heterozygote (barring *de novo* events). Using these sites only we can estimate:

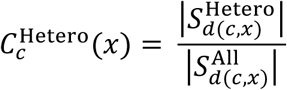

where *d*(*c*, *x*) is the depth at *x* for the child *c* and 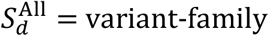 pairs (*f*′, *x*′) where the child *c*′ has depth *d*(*c*, *x*) = *d* at variant *x*′ and one of the parents is HomRef from the variant and the other parent is HomAlt after applying the sites filter and a conservative filter on the genotype quality of the parents, 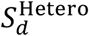 = the subset of 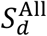 where the child is called as heterozygous and passes the heterozygote filter.

Similarly we can calculate:

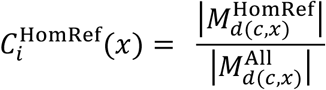

Where *i* is either *m* or *p* and 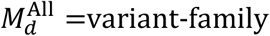 pairs (*f*′, *x*′) where all the children *c*′ has depth *d*(*c*, *x*) = *d*, and both parents in each family in question are HomRef for the variant, the variant is present in at least one of the other families after applying the sites filter and a conservative filter on the genotype quality of the parents.

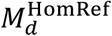 = the subset of 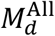 where the children are called as homozygous for the reference allele and passes the homozygote filter.

### Minimizing false positives *de novo* mutation calls

While the estimation of callability, as described above, reduces the effect of false negatives on the estimated mutation rate, it is still necessary to set the cutoffs in the filters so high that only very few or no false positives get into the set of estimated de novo mutations. We can fit the filter criteria by looking at the effect of different criteria on the rate estimate and the effect on how large a fraction of the called *de novo* variants are present in unrelated individuals from the same species. (Supplementary Fig. S1).

Based on these considerations we set the filter values at:

- *GQ ≥ 60* (for both the homozygote and heterozygote filter)
- *DP ≥ 10* (for both the homozygote and heterozygote filter)
- *RC <1.9* (for both the homozygote and heterozygote filter)
- *AD2 = 0* (for the homozygote filter)
- *lowQ_AD2 = 1*
- *AlleleBalance* > 0.3
- *minStrandCount* = 1

The AlleleBalance filter was set based on the distribution of AlleleBalance in the children after applying the other filters

### Parent of origin assignment of *de novo* mutations

We estimate the paternal origin of each *de novo* variant using the same strategy used in Maretty et al. (*1*)

For each variant, X, we use o(X) to denote the parental origin of the alternative allele. The reads might provide conflicting evidence and to find the most likely parental origin we calculated a likelihood ratio comparing how likely it is that the alternative allele is on the paternal chromosome (o(X)=1) to how likely it is that the alternative allele is on the maternal chromosome (o(X)=0):

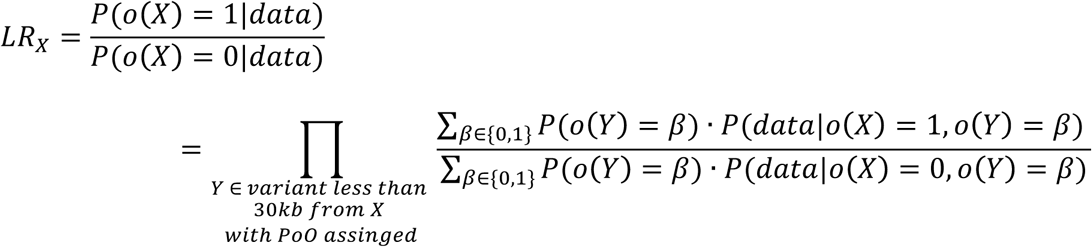

If *LR*_*X*_ is above one it indicates that the alternative allele of variant X is on the paternal chromosome and if *LR*_*X*_ is below one it indicates that it is on the maternal chromosome. The data that is informative about the PoO are the reads that cover both X and Y:

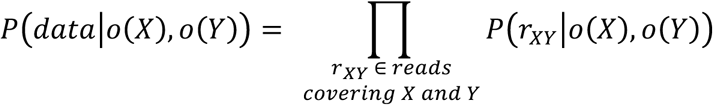

The probability that a read supports the true phasing is 1 if the read is mapped correctly and ½ if the read is not mapped correctly. We calculated the conditional probability of the read as:

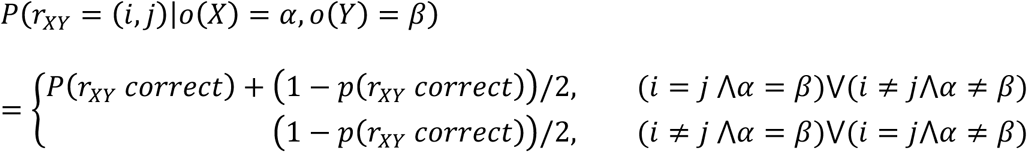

where *P*(*r*_*XY*_ correct) is the probability that *r*_*XY*_ is mapped correctly (estimated from the phred score in the bam file) and the values of *i* and *j* is either “ref” or “alt” depending on whether the read contains the reference allele or the alternative allele at position *X* and *Y*. For inherited variants where the parental origin could be assigned by just looking at the genotypes of the family members, *p*(*o*(*Y*) = 1) is calculated using the phred-scaled genotype probabilities of the three family members. If the PoO of variant Y has been assigned using read information we calculate *p*(*o*(*Y*) = 1) from the estimated LR: *p*(*o*(*Y*) = 1) = *LR*_*Y*_/(*LR*_*Y*_ + 1) The assignment of parental origin is carried out iteratively until no additional variants can be assigned.

### Estimation of yearly mutation rate

The mutation rate estimates for each of the great apes trios were converted into yearly estimates by extrapolating from the relationship between mutation rate and maternal and paternal parent age observed in humans. In humans, the best estimate is that the mutation rate in a child depend on the parental ages as follows:

Mutation rate = 1.77E-09 + 7.26E-11 * maternal age + 2.87E-10 * paternal age.

From this relationship, we calculate the expected mutation rate for each great ape trio taking parental ages into account and assuming the overall mutation rate per year is as in humans. The relative mutation rate for each trio is then calculated as the observed rate divided by the expected rate from the relationship in humans with 95% confidence intervals.

### Phylogenetic dating

We produced estimates of genomic divergence rates using the relative rates observed in each trio in the following way; from Scally et al. (*10*) we obtained the following average genomic divergences between:

Human and chimpanzee = 0.0137

Human and gorilla = 0.0175

Human and orangutan = 0.034

The great apes phylogeny deviates slightly from a molecular clock, according to Moorjani et al.

(*7*) with the chimpanzee branch being 2% longer than the human branch, the gorilla branch 6% longer than the human branch since their common ancestry and the orangutan branch 11% longer than the human branch since their common ancestry. Using these numbers and focusing on the human branch the branch lengths from human to the common ancestor with the chimpanzee becomes 0.006713, with gorilla 0.008225 and with orangutan 0.01513.

Using the estimated chimpanzee mutation rate (0.72 per billion years) since the common ancestry between human and chimpanzee this corresponds to 9.46 my for human-chimpanzee average genomic divergence time, using the gorilla rate (0.66) since the common ancestry between human and gorilla this becomes 12.42 million years for human-gorilla average genomic divergence time, and using the orangutan rate (0.00079) since their common ancestry becomes 21.85 million years for the average human-orangutan geomic divergence time.

To turn the divergence numbers into estimates of species separation time (here equalled to speciation time) we used the ancestral effective population sizes reported in Scally et al. (*10*) scaled to the mutation rates assumed in the common ancestors yielding:

Human chimpanzee ancestral effective population size = 70,125

Human gorilla ancestral effective population size = 74,361

Human orangutan ancestral effective population size = 127,213

Since the expected coalescence time in the common ancestors are 2*N*, the separation times are calculated as:

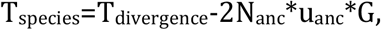

Where G is the generation time, here assumed to be 25 years (approximate average of generation times in extant species, humans 29, chimp 24, gorilla 19, orangutan 25)

This yields the following estimates

Human chimpanzee speciation time = 5.96 million years

Human gorilla speciation time = 8.70 million years

Human orangutan speciation time = 15.49 million years

Numbers are summarised in Figure 3.

## Acknowledgments

We thank Molly Przeworski and Priya Moorjani for comments to a previous version of the manuscript.

## Author contribution

Designed the study: S.B., T.M., T.M.-B, C.H., M.H.S. Contributed reagents: C.H. Performed analysis: S.B., T.M., M.H.S. Wrote the paper: S.B., M.H.S. with input from all authors.

## Data and materials availability

All sequence data will be submitted to EGA.

---

**Figure S1.**
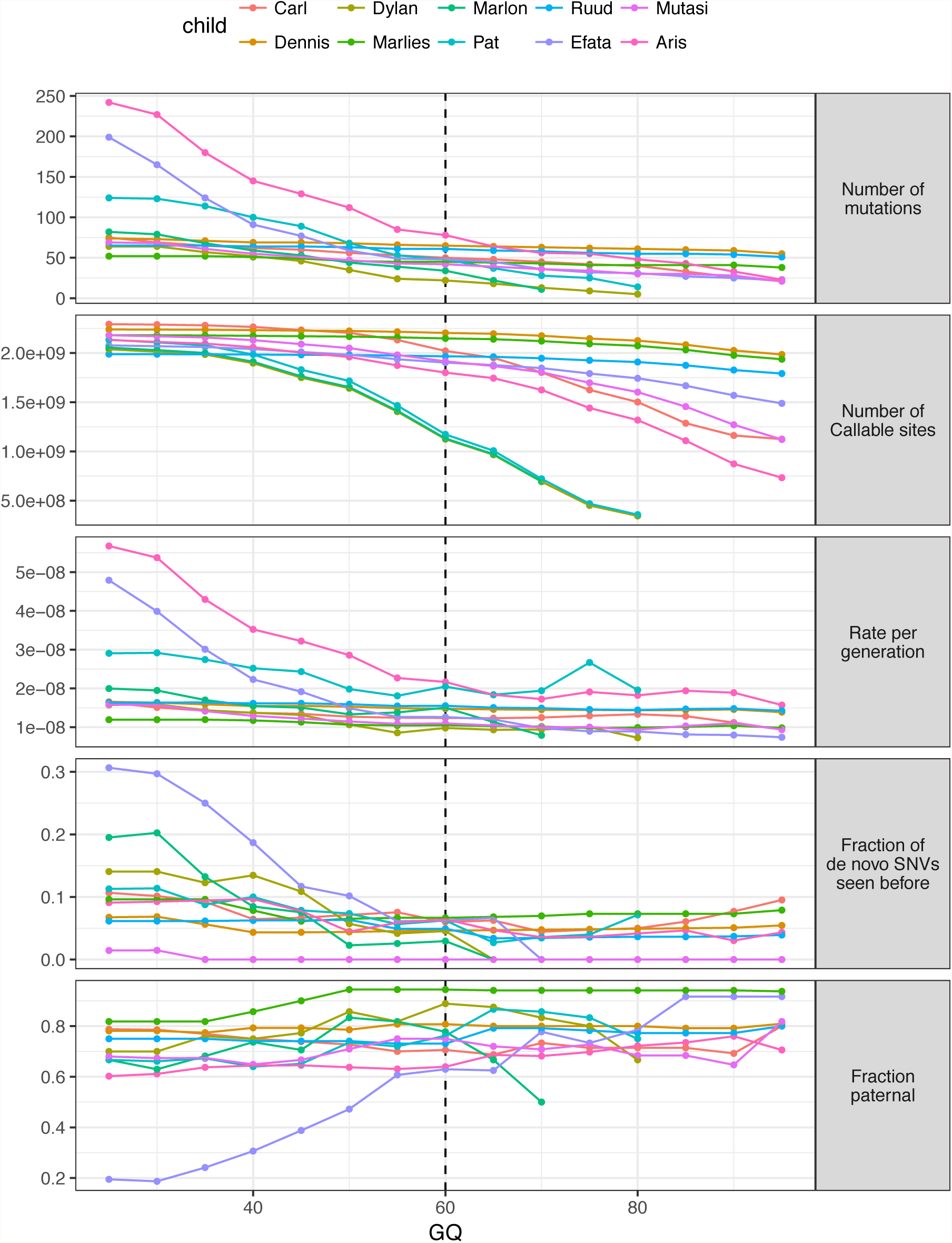
Determining quality threshold of genotype quality in the families. For each quality threshold (GQ), the Figure shows a. Number of mutations called, the callable number of genomic baspairs for that quality, c. these two divided as the mutation rate for the given child, d. the fraction of the called de novo mutations that are annotated as SNVs in databases, and e. the fraction of the mutations that can be assigned to the paternal parent (given that they can be assigned to one of the parents, typically 70% of the mutations, see Table 1)

**Figure S2.**
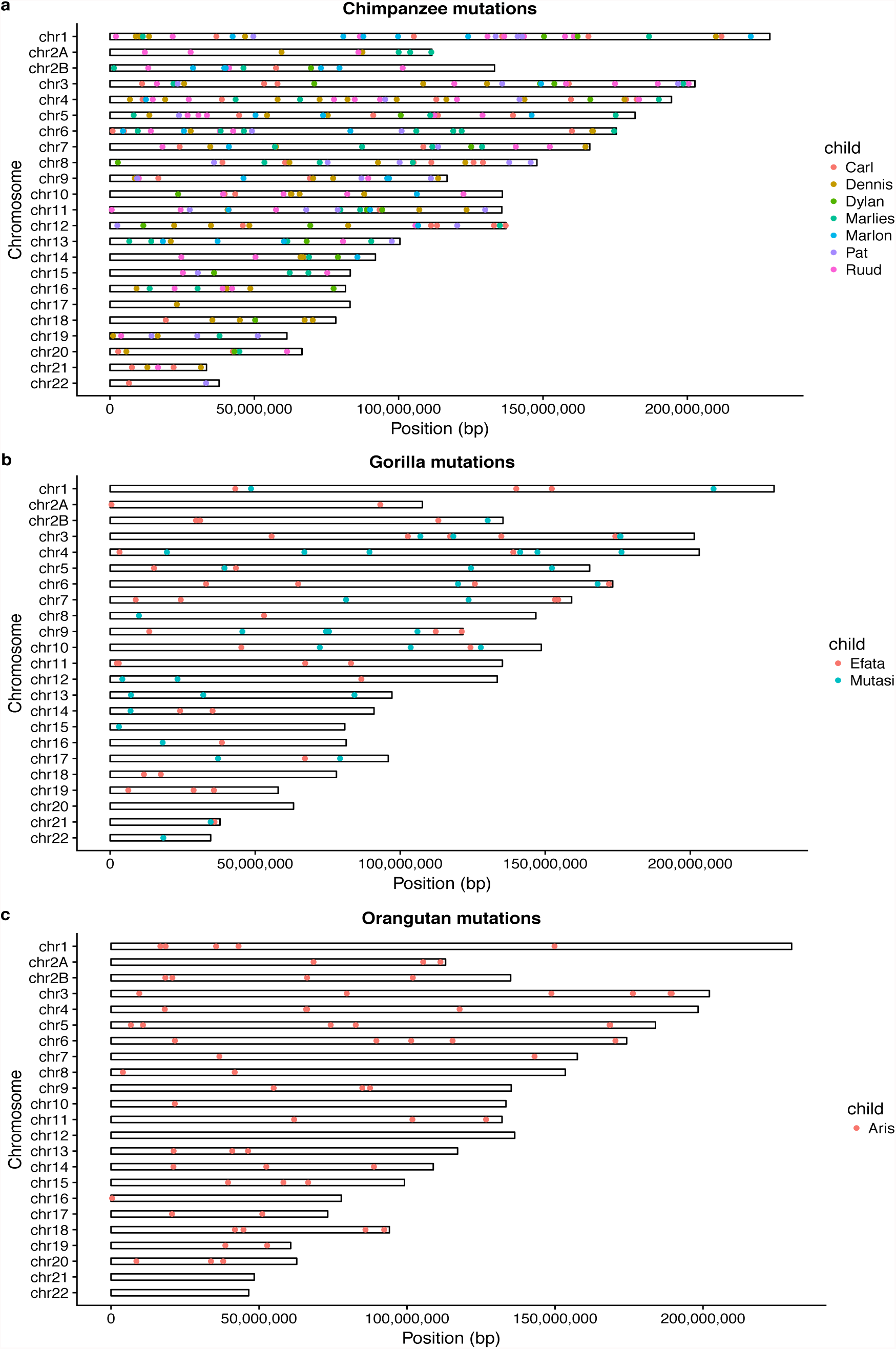
The position of the called de novo mutation along the different reference genomes of the great apes. a. Chimpanzee, b. Gorilla, c. Orangutan

**Figure S3.**
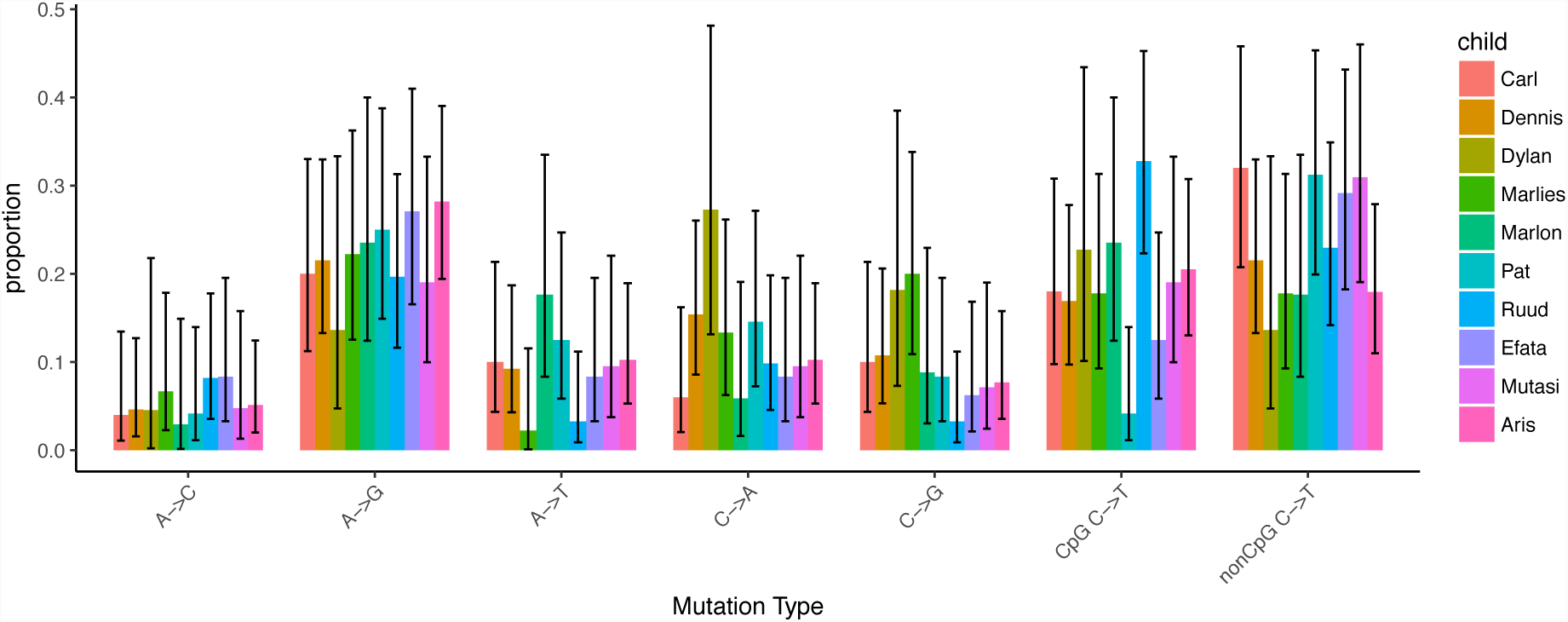
The proportion of different types of mutations in the ten trios (95% CI). The C→T mutations are divided into the proportion in CpG context and the proportion not in CpG context.

**Figure S4.**
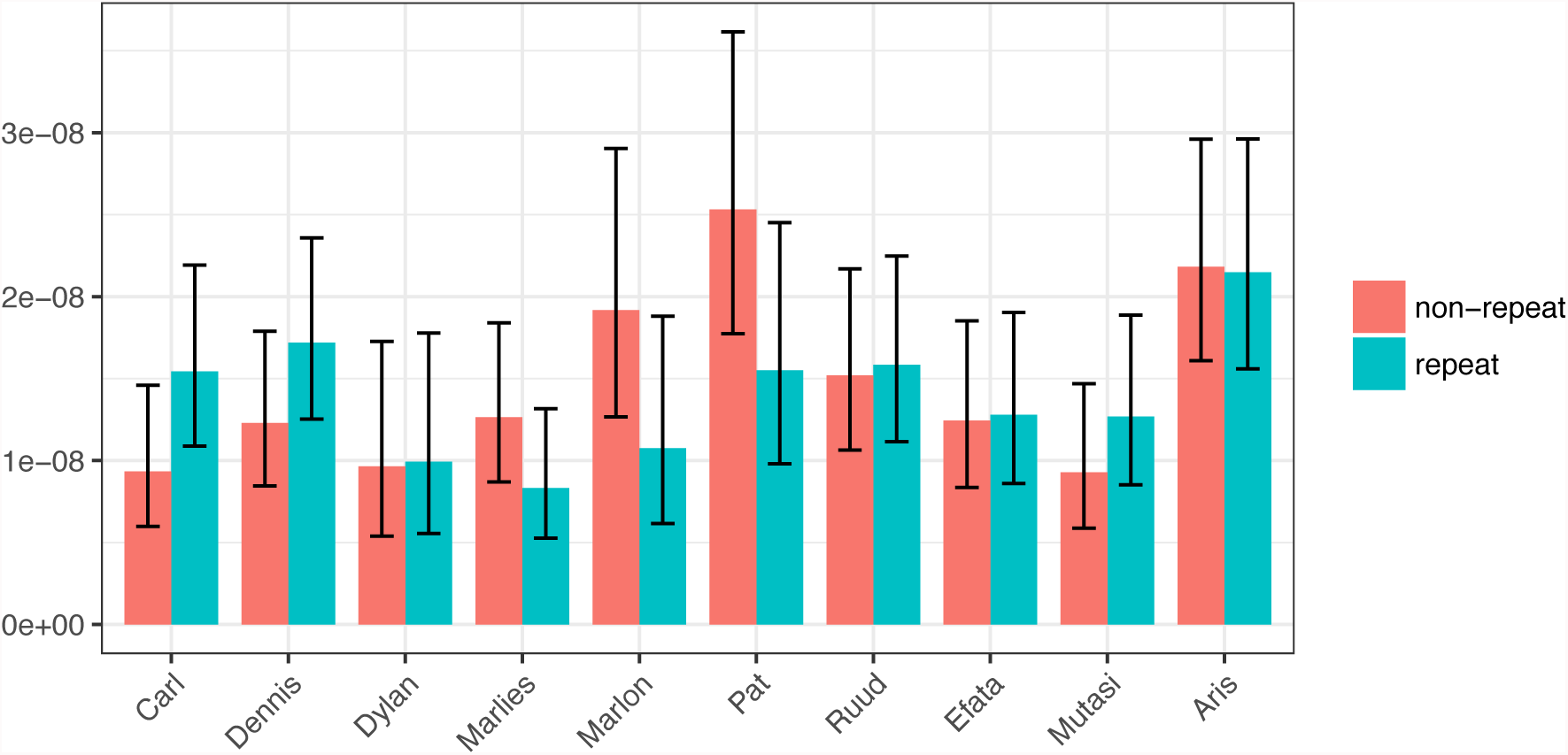
The mutation rate estimated for the non-repetitive and repetitive (from repeatmaker) part of the reference genomes (95% CI). The numbers can be found in Table S1 and S2. In none of the individuals are there significant differences between the regions.

**Figure S5.**
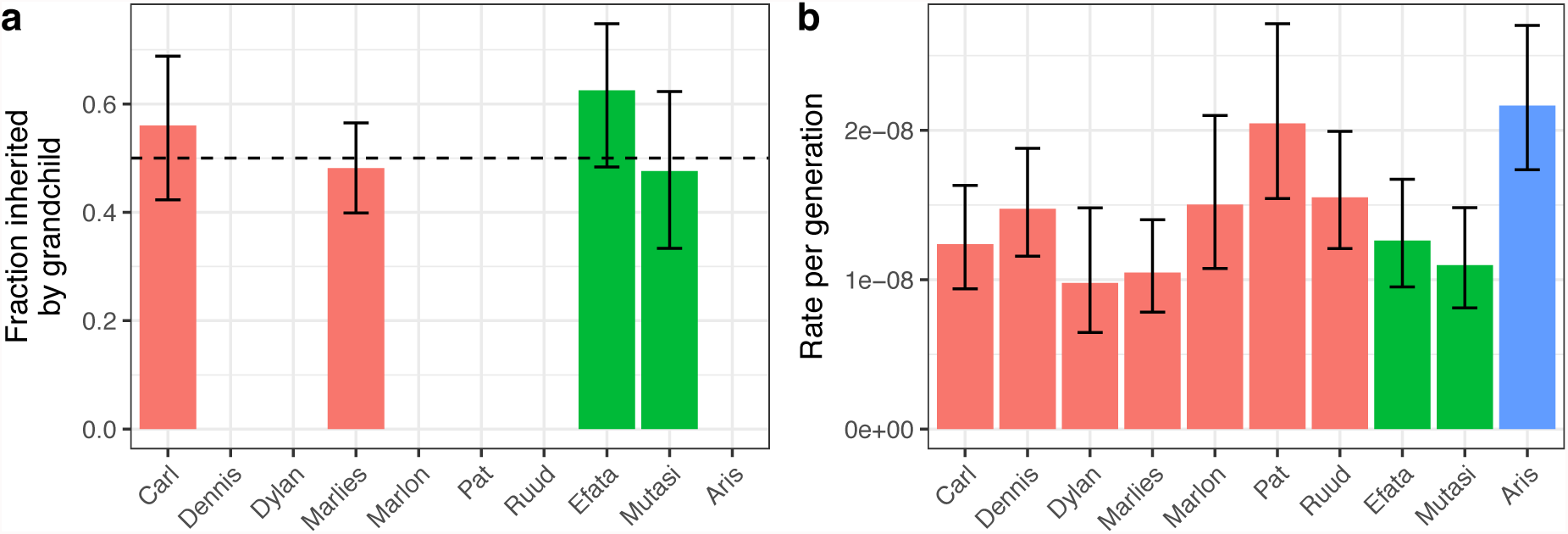
Comparison of key mutational parameters for the different trios. a) The fraction of mutations inherited by grandchild when available (95% CI). b) The rate per generation (95% CI)

**Figure S6.**
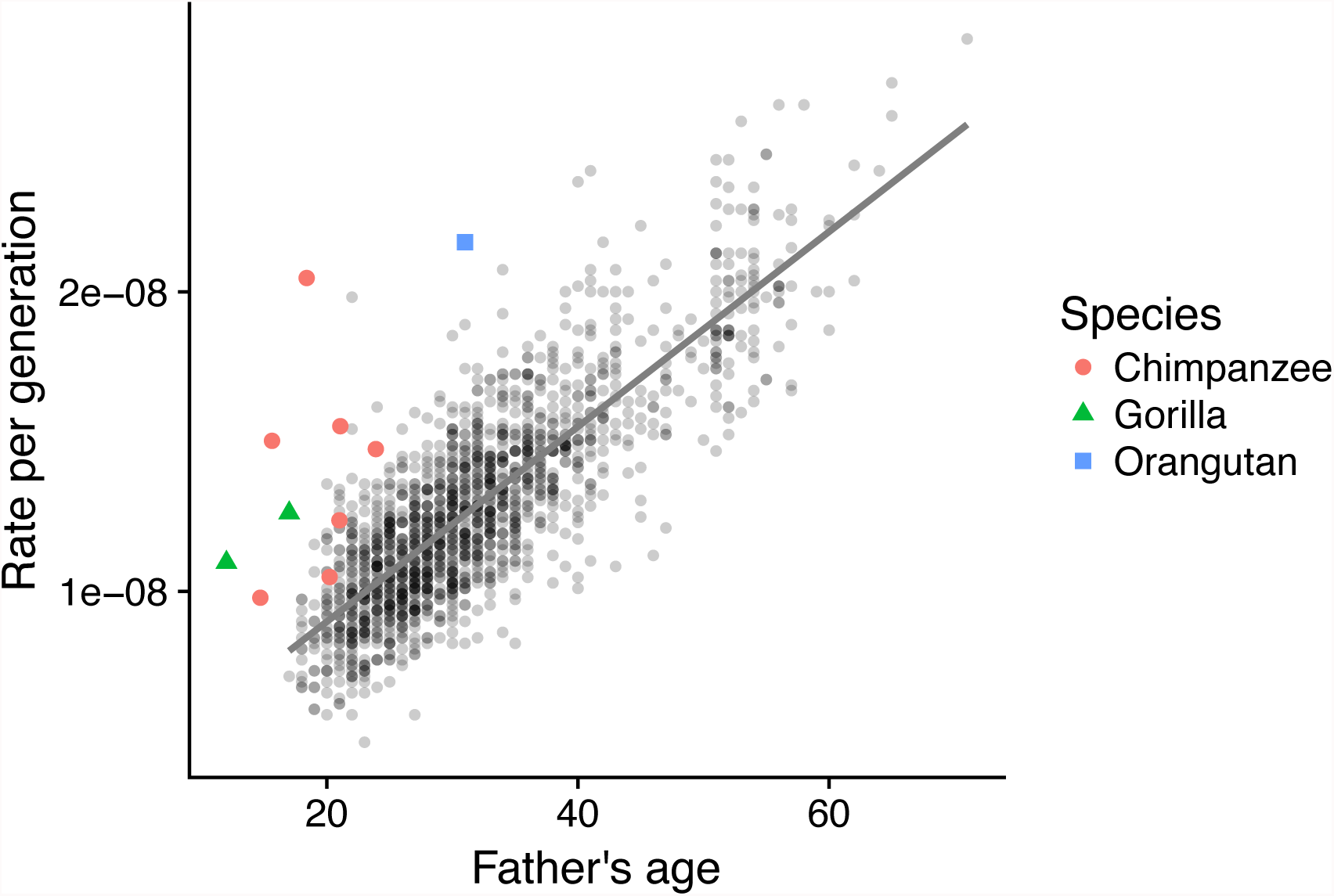
The estimated mutation rate as a function of paternal age for 1536 human trios (grey point, with linear regression) and for the great apes trios investigated in the present study

**Figure S7.**
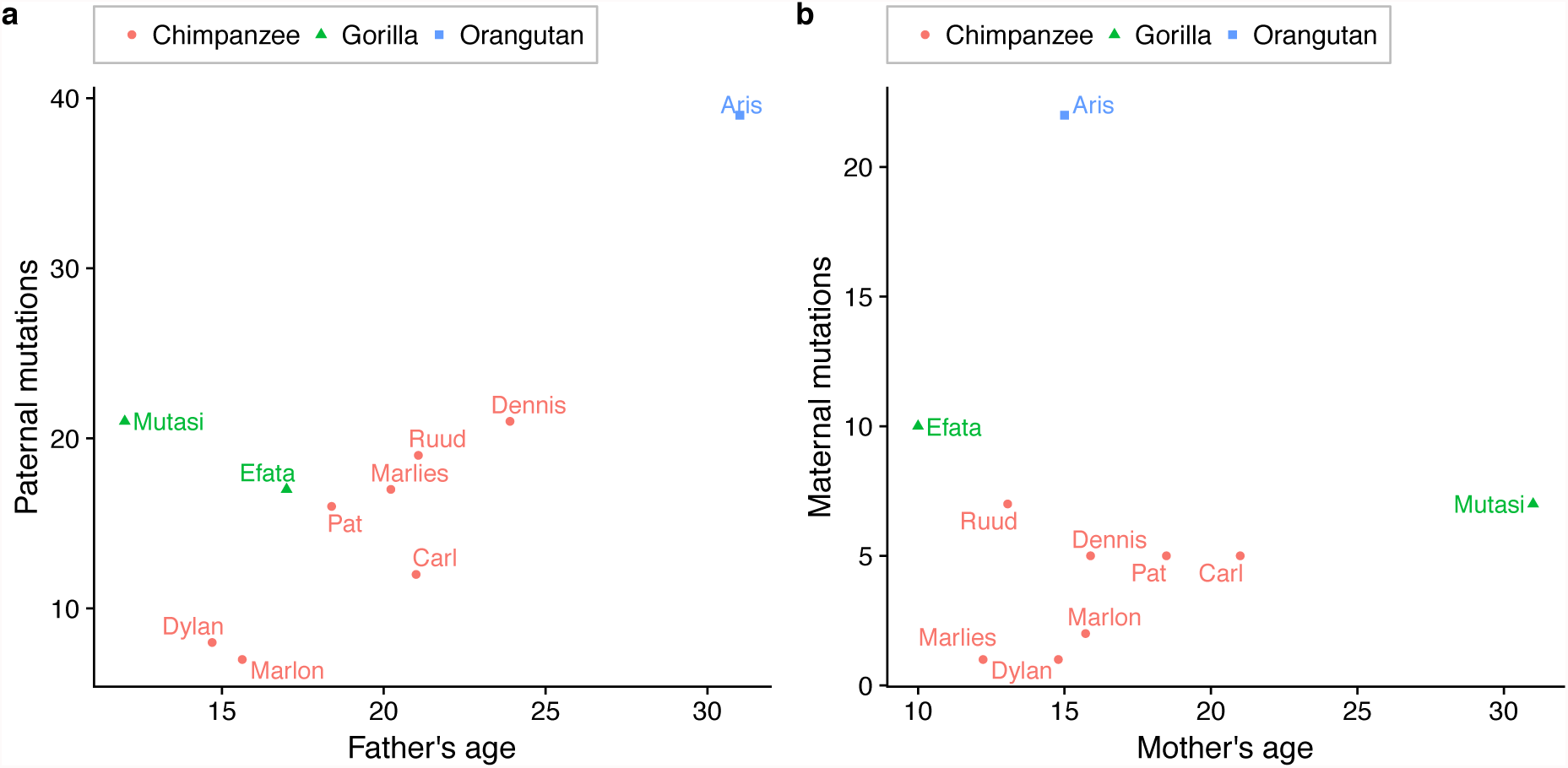
The number of mutations passed on from the parents as a function of their age. a. paternal mutations as function of fathers age, b. maternal mutations as function of mothers age

**Supplemental Table S1:** Summary data for *de novo* SNV counts of different types

| Individual<br>(Species) | Carl<br>(Chimpanzee) | Dennis<br>(Chimpanzee) | Dylan<br>(Chimpanzee) | Marlies<br>(Chimpanzee) | Marlon<br>(Chimpanzee) |
| --- | --- | --- | --- | --- | --- |
| all mutations | 50 | 65 | 22 | 45 | 34 |
| CpG | 9 | 12 | 6 | 9 | 8 |
| CpG transitions | 9 | 11 | 5 | 8 | 8 |
| non-CpG Strong (C or G) | 24 | 30 | 12 | 22 | 11 |
| Weak (A or T) | 17 | 23 | 4 | 14 | 15 |
| transitions | 35 | 39 | 11 | 26 | 22 |
| transversions | 15 | 26 | 11 | 19 | 12 |
| Previously observed | 3 | 3 | 1 | 3 | 1 |
| Paternal | 12 | 21 | 8 | 17 | 7 |
| Maternal | 5 | 5 | 1 | 1 | 2 |
| In a repeatmasked region | 31 | 38 | 11 | 18 | 12 |
| outside repeatmasked<br>regions | 19 | 27 | 11 | 27 | 22 |

| Individual<br>(Species) | Pat<br>(Chimpanzee) | Ruud<br>(Chimpanzee) | Efata<br>(Gorilla) | Mutasi<br>(Gorilla) | Aris<br>(Orangutan) |
| --- | --- | --- | --- | --- | --- |
| all mutations | 48 | 61 | 48 | 42 | 78 |
| CpG | 3 | 20 | 7 | 8 | 19 |
| CpG transitions | 2 | 20 | 6 | 8 | 16 |
| non-CpG Strong (C or G) | 25 | 22 | 20 | 20 | 25 |
| Weak (A or T) | 20 | 19 | 21 | 14 | 34 |
| transitions | 29 | 46 | 33 | 29 | 52 |
| transversions | 19 | 15 | 15 | 13 | 26 |
| Previously observed | 3 | 3 | 3 | 0 | 5 |
| Paternal | 16 | 19 | 17 | 21 | 39 |
| Maternal | 5 | 7 | 10 | 7 | 22 |
| In a repeatmasked region | 18 | 31 | 24 | 24 | 37 |
| outside repeatmasked<br>regions | 30 | 30 | 24 | 18 | 41 |

**Supplemental Table S2:** The number of callable sites for different types of base pairs. With numbers from Supplementary Table S1, these can be used to estimate the mutation rates for different types of mutations

| Individual | Carl | Dennis | Dylan | Marlies | Marlon |
| --- | --- | --- | --- | --- | --- |
| (Specie) | (Chimpanzee) | (Chimpanzee) | (Chimpanzee) | (Chimpanzee) | (Chimpanzee) |
| all sites | 4041209937 | 4406543730 | 2248382773 | 4295124822 | 2262663575 |
| CpG | 65502956 | 79235211 | 30524383 | 78187818 | 30688052 |
| non-CpG String (C or G) | 1521027591 | 1702394038 | 810469237 | 1665488365 | 815549823 |
| Weak (A or T) | 2454679389 | 2624914481 | 1407389153 | 2551448639 | 1416425701 |
| In a repeatmasked region | 2007047695 | 2210655941 | 1107561174 | 2160655070 | 1115357595 |
| outside repeatmasked regions | 2034162242 | 2195887789 | 1140821599 | 2134469752 | 1147305980 |

| Individual | Pat | Ruud | Efata | Mutasi | Aris |
| --- | --- | --- | --- | --- | --- |
| (Specie) | (Chimpanzee) | (Chimpanzee) | (Gorilla) | (Gorilla) | (Orangutan) |
| all sites | 2345308475 | 3931520576 | 3804369957 | 3830238966 | 3600614878 |
| CpG | 32060877 | 69951969 | 57718100 | 62181326 | 67571766 |
| non-CpG String (C or G) | 847254888 | 1516602242 | 1436039179 | 1454519685 | 1389697054 |
| Weak (A or T) | 1465992710 | 2344966365 | 2310612678 | 2313537955 | 2143346058 |
| In a repeatmasked region | 1160691764 | 1957168353 | 1876167556 | 1892445281 | 1721785714 |
| outside repeatmasked regions | 1184616711 | 1974352223 | 1928202400 | 1937793685 | 1878829164 |

